# Combining biomarker and virus phylogenetic models improves epidemiological source identification

**DOI:** 10.1101/2021.12.13.472340

**Authors:** Erik Lundgren, Ethan Romero-Severson, Jan Albert, Thomas Leitner

## Abstract

To identify and stop active HIV transmission chains new epidemiological techniques are needed. Here, we describe the development of a multi-biomarker augmentation to phylogenetic inference of the underlying transmission history in a local population. HIV biomarkers are measurable biological quantities that have some relationship to the amount of time someone has been infected with HIV. To train our model, we used five biomarkers based on real data from serological assays, HIV sequence data, and target cell counts in longitudinally followed, untreated patients with known infection times. The biomarkers were modeled with a mixed effects framework to allow for patient specific variation and general trends, and fit to patient data using Markov Chain Monte Carlo (MCMC) methods. Subsequently, the density of the unobserved infection time conditional on observed biomarkers were obtained by integrating out the random effects from the model fit. This probabilistic information about infection times was incorporated into the likelihood function for the transmission history and phylogenetic tree reconstruction, informed by the HIV sequence data. To critically test our methodology, we developed a coalescent-based simulation framework that generates phylogenies and biomarkers given a specific or general transmission history. Testing on many epidemiological scenarios showed that biomarker augmented phylogenetics can reach 90% accuracy under idealized situations. Under realistic within-host HIV evolution, involving substantial within-host diversification and frequent transmission of multiple lineages, the average accuracy was at about 50% in transmission clusters involving 5-50 hosts. Realistic biomarker data added on average 16 percentage points over using the phylogeny alone. Using more biomarkers improved the performance. Shorter temporal spacing between transmission events and increased transmission heterogeneity reduced reconstruction accuracy, but larger clusters were not harder to get right. More sequence data per infected host also improved accuracy. We show that the method is robust to incomplete sampling, and we evaluate real HIV-1 transmission clusters. The technology presented here could allow for better prevention programs by providing data for locally informed and tailored strategies.

**AUTHOR SUMMARY:** While phylogenetic methods have been successfully used to infer transmission patterns on many scales, substantial uncertainty about the corresponding transmission history has remained. Here, we introduce a formal inference framework for the simultaneous use of HIV biomarkers and HIV sequences derived from persons who may have infected each other. We created a flexible system that can use up to five biomarkers to estimate the time of infection of each person, with the appropriate uncertainty given by an empirical probability distribution. These time-of-infection distributions were jointly modelled with the HIV phylogenetic tree estimation to produce possible transmission histories. We show that adding biomarkers substantially limits the possible transmission histories. This makes it possible to identify transmission risks, to assess confidence in source attributions, and to make efficient resource allocations to prevent further transmissions in local epidemics.

## INTRODUCTION

To effectively control an infectious disease, limited prevention resources must be allocated to where they are needed most [1]. Thus, identifying hotspots of transmission would allow for efficient resource allocation. Transmission, especially in chronic infections such as HIV, is heterogenous in time, place, and person, leading to episodic transmissions and local outbreaks. Mapping these events using traditional epidemiological methods is challenging, expensive, and slow, and may be inaccurate. For example, several studies have reported that interview-based information about sexual contacts where HIV transmission might have taken place was often not in agreement with the phylogenetic history of the transmitted virus [2, 3]. Therefore, phylogenetic reconstruction, using existing and growing public health databases, provides an attractive alternative.

Reconstructing the history of an epidemic using phylogenetic methods has become a substantial domain of phylodynamic research involving many different pathogens [4–11]. The primary technical challenge in this domain is that the typical mode in which genetic sequences are sampled, either in terms of a study or through a surveillance system, may not be sufficient for reconstructing transmission histories. This applies especially for chronic infections where the pathogen develops substantial within-host diversity, such as in HIV, HBV, HCV, and some bacterial infections [12–16]. This within-host diversity means that when a person infects another, there are many alternative phylogenetic lineages that could have been involved, often more than one, leading to a non-trivial and non-identical correspondence between the transmission history and the pathogen phylogeny [12, 17, 18]. The extent of this problem was quantified by Hall and Coljin’s method that counts the exact number of transmission histories that are logically consistent with a pathogen phylogeny [19]. For example, a phylogeny from 20 infected persons (with 1 sequence/person) could have as many as 102 million transmission histories that are consistent with that phylogeny—the exact number depends on the observed phylogenetic topology. While additional constraints and Bayesian inference can overcome weak non-identifiability, it is desirable to have constraints that are both measurable (i.e., empirical) and based on readily available data. While the basic theoretical underpinnings of how the order of infection events constrains the set of possible transmission histories for a given phylogeny have been understood for several years [20], this knowledge currently is not integrated into phylodynamic inference methods. Some software such as SCOTTI [7] are able to include user-defined infection windows, i.e., a fixed period of time that a host was contagious. Unfortunately, for the case of life-long, chronic infections such as HIV, in the absence of treatment such a window would be too long to be useful, and typically the actual start of contagiousness is rarely known.

HIV biomarkers offer an alternative to infection windows, instead estimating when a host was infected [21]. Here, we introduce the use of HIV biomarkers to augment phylogenetic reconstruction and narrow down the possible transmission histories among epidemiologically linked hosts. Some such biomarkers are always available in clinical and public health databases, including HIV *pol* sequences, CD4 cell counts, and viral load measurements, and sometimes quantitative serological assay test results. In addition, there may be information about previous negative HIV test results and other demographic information that may limit possible time of infection. We show that it is possible to enhance transmission history reconstruction by modeling multiple biomarkers in a joint biomarker-phylogeny-transmission history framework.

## MATERIALS AND METHODS

### Methodological Overview

A transmission history is defined as who infected whom and when those transmission(s) occurred. While HIV transmissions can occur in many different contact networks [22], in this work we will consider 5-50 individual hosts that have transmitted HIV in different time intervals. When attempting to reconstruct a local transmission history, one can divide the available information into three levels, information from the sampling times, information from the genetic sequences, and information about the infection times (Fig 1). If only the sampling times were known, then the virus phylogeny and transmission history would be almost completely unconstrained. Adding sequence data limits the set of possible transmission histories by revealing temporal evolutionary relationships among sampled pathogens, which constrains the transmission history to some extent, but still leaves a wide variety of possibilities. With the additional information about the host’s infection times, the transmission tree is still not fully constrained, but there are many fewer possibilities than with sequences alone. In the example in Fig 1 (right panel), we can be quite certain that the red individual is the first infection in the cluster, but the order of the other two infections is not certain. In this particular case, there are nine possible transmissions among three persons, out of which all are plausible given the phylogeny. (There are 12 possible transmission histories if one includes the order of transmissions if one host infects both others, while only considering who-infected-whom in whatever order gives 9 possible transmission directions.) When including the probability density functions for the infection times, however, the number of plausible transmission trees is reduced to three. On the other hand, the phylogeny does constrain the infection times somewhat; for example, the red and green individuals could not have both been infected before their coalescence time.

**Fig 1:**
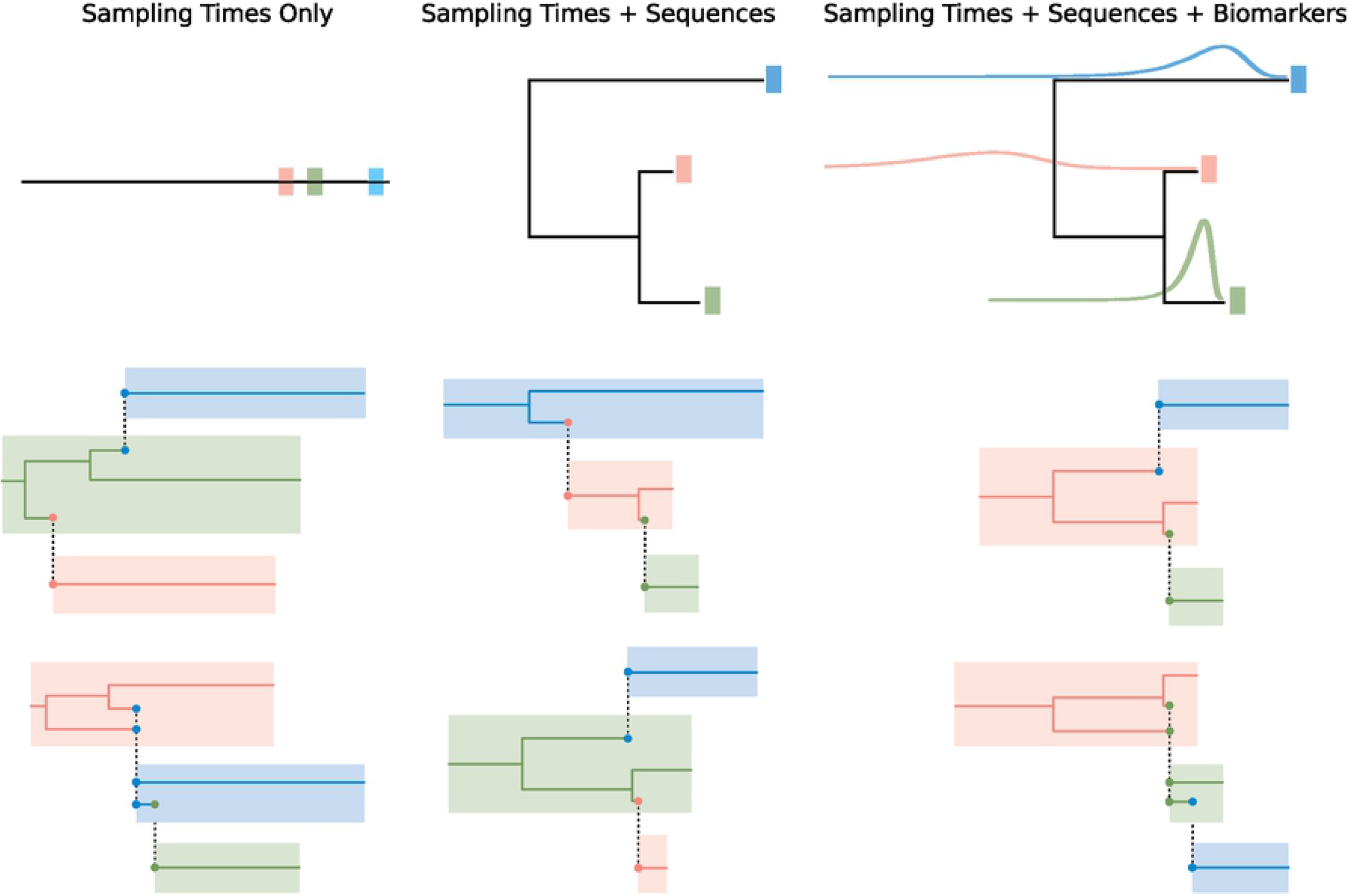
Conceptual Model Motivation. The level of information available for inference (top row) limits the possible transmissions between hosts (colored boxes). The bottom two rows show examples of inferred transmission histories given the different levels of information, with transmissions indicated by dashed vertical lines, and phylogenetic trees. The left side of each colored box in the phylogeny represents the time of infection, and the horizontal lines are the lineages present in each individual over the course of the outbreak, forming the phylogeny. Biomarkers are shown as probability densities of time of infection of each host. With all information levels available, transmission history inference becomes more constrained and thus improved.

### Multiple Biomarker Model

HIV biomarkers are measurable biological quantities that have some relationship to the amount of time someone has been infected with HIV. Based on the values of a set of biomarkers, we can infer a probability distribution for the amount of time that that person has been infected using a mixed effects Multiple Biomarker Model (MBM). Our model extends the previous model by Giardina et al 2019 [21], adding two additional biomarkers (bringing the total to five) and allowing inference when not all biomarker measurements are available. We also scaled the biomarker values such that the input values to the model are all at the same order of magnitude and used a prior distribution that more closely reflects the expected amount of time between infection and diagnosis.

The mixed effects model is of the form 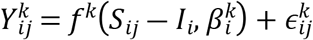, where 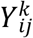 is the measured value of the *k*^th^ biomarker at the *j*^th^ timepoint for the *i*^th^ individual, *f*^*k* is the function that predicts the value of a biomarker based on time after infection, *S_ij_* is the time of the *j*^th^ sample for the *i*^th^ individual, *I_i_* is the infection time of the *i*^th^ individual, 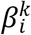 are the function parameters for the *k*^th^ biomarker for the *i*^th^ individual, and 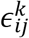 is the biological and measurement noise.

We modeled five biomarkers: BED, the IgG capture BED enzyme immunoassay [23]; LAg, Limiting Antigen Avidity assay [24]; *pol* polymorphism count, the number of multi-state nucleotide characters in *pol* direct population sequences [25, 26]; *pol* NGS diversity, HIV diversity estimated from next-generation sequencing of *pol* on the Illumina platform [27]; and CD4 cell count, the number of CD4 positive T cells in 1 ml plasma. As shown in Fig 2, BED and LAg are modeled as log10 values, normalized with an internal test standard, starting at low concentrations that rise asymptotically over time since infection as in Skar et al [28], with LAg typically rising faster than BED. Both the *pol* polymorphism count and the *pol* NGS diversity (of 3rd codon positions) increase approximately linearly at the timescales that we are interested in. The CD4 cell count is modelled as the square root in order to make the time series trend more linear.

**Fig 2:**
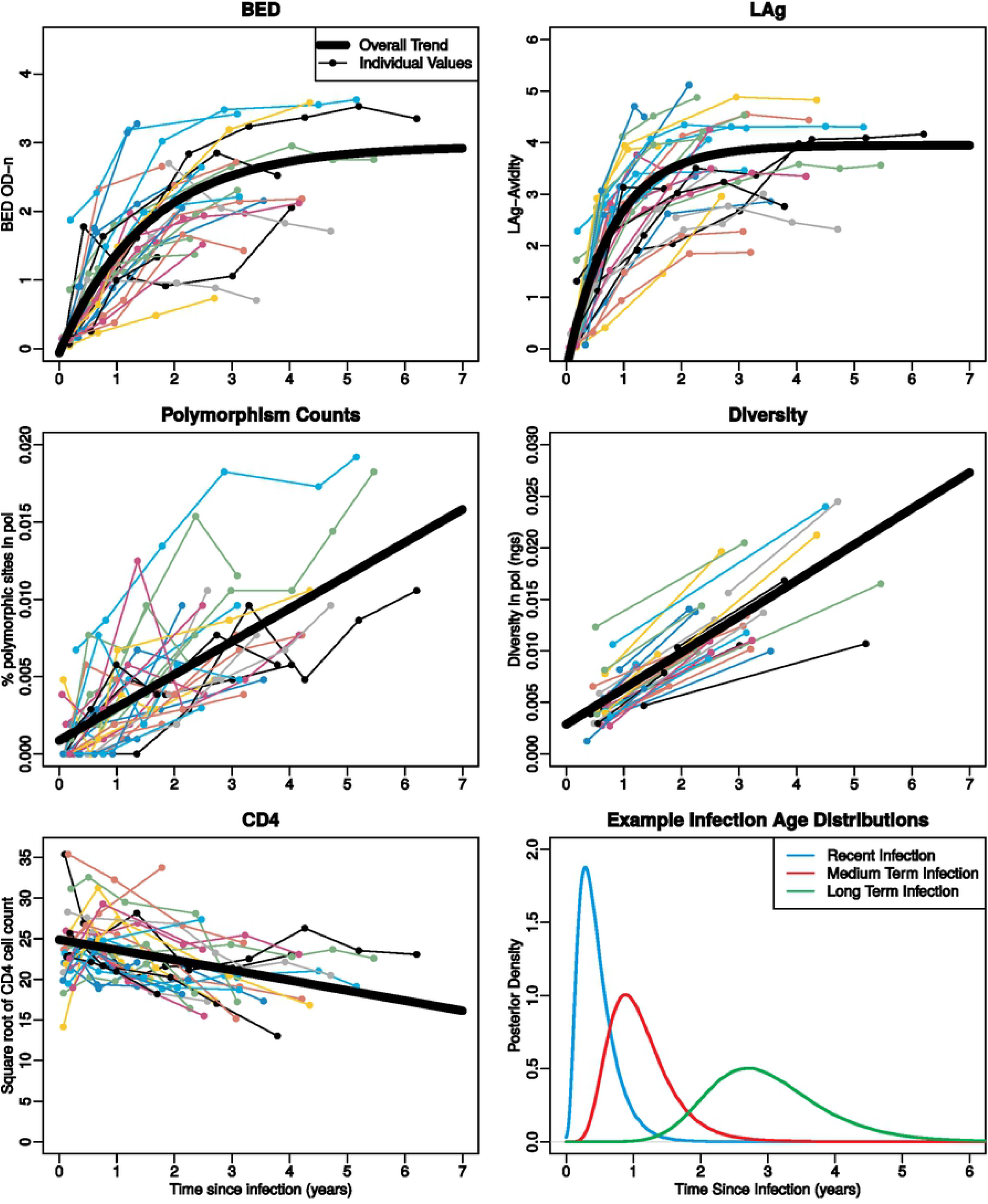
HIV Biomarker time trends. Time series plots for each of the five biomarkers used in our modeling for a group of 30 patients (colored lines). The best fit line for the fixed part of our mixed effects model is shown for each biomarker (bold black lines). The bottom right panel shows examples of the inferred distributions of infection age using biomarker values from a recently infected patient, a patient who has been infected somewhat longer, and a longer-term infected patient.

We used a Markov Chain Monte Carlo (MCMC) sampler implemented in *rjags* [29] to sample from the posterior distributions of the infection times. The prior distribution for the infection times was a Gamma distribution with mean 2 years and standard deviation 1.5 years, chosen to be qualitatively similar to the distributions for time between infection and diagnosis found in Giardina et al. 2019 [21]. We created a flexible system that can use any number or combination of biomarkers (including no biomarkers, which would recover the prior distribution).

### Biomarker Training Data and Validation

The training data for our multiple biomarker model came from 30 longitudinally followed Swedish HIV-infected persons with well-defined times of infection. Biomarker data from these patients have been previously used: *pol* polymorphism, BED, and CD4 counts were used in Giardina et al [21]; and *pol* NGS diversity in Puller et al [27]. In this study we added LAg for a total of five biomarkers. The 30 patients were selected to have: 1) a previous negative test and first positive test that were no more than 6 months apart or a known primary HIV infection time, 2) at least three follow up measurements over a long time period (2-5 years), and 3) been treatment naïve for that time period. The biomarker data were measured on stored biobank samples because the inclusion criteria are difficult to fulfill as modern clinical practice is to put patients on antiviral treatment immediately after HIV diagnosis. The full model was trained using all measurements from all 30 patients, using 5×10^4^ iterations of model adaptation, 3×10^5^ iterations of burn-in, and 1×10^6^ samples.

To validate our MBM, we performed a leave one out cross validation using one set of biomarker measurements from each of the 30 patients as testing data while using all measurements from the other 29 patients as training data. The MBM was also used to simulate realistic biomarker values for testing purposes: We first found the maximum likelihood values of the model parameters (trained on all 30 patients), then used those values to simulate new random effects trajectories for the expected values of each of the biomarkers over time, and, finally, simulated new biomarker values by adding Gaussian noise to the expected values.

### Joint Inference of Transmission History and Phylogeny

The MBM-derived posterior distributions of the infection times were incorporated into the likelihood function for the transmission history and phylogenetic tree. The general form of the likelihood function remains the same as in Klinkenberg et al 2017 [6]:

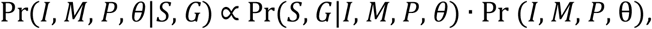

with unobserved infection times *I*, infectors *M*, phylogeny *P*, parameters *θ*, observed sampling times *S*, and genetic sequences *G*. This likelihood function can be split up into four terms for the likelihood of the sequences, the phylogenetic tree, the difference between the infection and sampling times, and the transmission tree as well as a term for the prior distributions of the parameters:

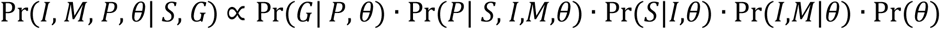

Given the distributions, we performed the transmission history and phylogenetic inference using a modified version of the R package *phybreak* [6]. We refer to our modified version as *biophybreak* (https://github.com/MolEvolEpid/biophybreak). We modified a version of *phybreak* updated by the original author, with the most notable change since the publication of Klinkenberg et al 2017 [6] being the inclusion of the possibility of a wide transmission bottleneck. This modification is important to model realistic HIV transmission where >1 phylogenetic lineage often is transmitted [30]. The primary modification that we introduced here (in *biophybreak*) is the way in which the likelihood for the interval between infection and sampling is calculated. In the original version of *phybreak*, the length of that interval is a Gamma distribution with a user specified shape and mean that is estimated as a model parameter, with the same parameters for every individual. Our modification allows any non-parametric distribution to be used for this likelihood as well as allow each individual to have their own distribution. Specifically, we used the posterior distribution of the infection age obtained from the MBM for each individual. Additionally, we added the Generalized Time Reversible (GTR) substitution model, which has been shown to be the most realistic HIV evolutionary model [31] (instead of the Jukes-Cantor model used in the original *phybreak* package). However, this comes at a computational cost of about four times longer MCMC iteration steps. The third modification we made allows a custom “generation function”, i.e., the function that specifies how likely an individual is to infect another individual based on how long they have been infected. This was motivated by the observation that a higher viral load, seen in the acute infection stage, results in a higher risk of transmission [32]. Since it is difficult to know exactly what the shape of the generation function should be, but it is known that approximately half of transmissions are from recently (within six months) infected individuals [33], we used a step function that is three times higher in the first six months, resulting in a fairly conservative improper prior distribution, which is fine in this case since only the relative values of the distribution are important. As with the infection time distributions, we allow any function to be used, e.g., modifying transmission probability before and after diagnosis, as described in the next section.

### General Simulation Experimental Design

In order to test our methodology and determine which types of transmission histories that may more or less difficult to correctly infer, we performed a variety of simulations. We varied the number of individuals, the frequency of infection events in the transmission history (temporal spacing), the heterogeneity of the number of transmissions per individual (standard deviation of the network degree), the level of information about infection time we get from the biomarkers, and the mutation model used in the simulation and inference (mutation parameters according to the HIV-1 *pol* or *env* gene).

For the simulation sets that use randomly generated transmission histories, we first specified the number of individuals, then the amount of time from the first to the last infection, taking into account the number of individuals and temporal spacing. The infection times were then generated using a continuous uniform random variable between the first and last infections, with the option to have a minimum amount of time between any two infection times, which we set to 0.05 years. Next, sampling times were given to each individual. Infectors (transmission donors) were chosen such that the resulting transmission tree had a transmission heterogeneity close to the desired value. To facilitate this, weights were assigned to each individual for how likely they were to be a donor, with the variation in weights depending on the target amount of transmission heterogeneity. The weights from the generation functions were also taken into account at this time. Note that since the infection times were chosen before the infectors rather than having the new infection times chosen from each infector’s generation function, the generation function was implicitly changed in a non-trivial way. Therefore, we also allowed *biophybreak* to use a penalty for transmission after diagnosis, which may be justified when all patients are successfully and continuously treated after diagnosis [34]. Hence, the weights of the potential infectors can be modified by a factor depending on whether they have been sampled yet. As in the case of the generation function, this post-sampling transmission penalty is implicitly changed by the way the infection times are chosen.

Given a transmission history, we created the phylogeny with a coalescence simulator that used a within-host model of linearly increasing effective population size *N*(*t*) = *α* + *βt*, where *α* is the effective population size at the time of infection and *β* is the growth rate of the effective population size per generation [17, 35], with a generation assumed to be 1.5 days [36]. Unless otherwise noted, we used *α* = 5 and *β* = 5. Finally, sequences were generated with SeqGen [37] using known absolute and relative substitution rate parameters from either the HIV-1 envelope gene (*env*) or polymerase gene (*pol*).

We performed the transmission history and phylogenic inference with *biophybreak* using 2×10^5^ MCMC samples after 5×10^4^ iterations of burn-in unless otherwise noted. We used an effective sample size (ESS) of 200 for the model parameters ensure proper mixing of the MCMC chains. We used two different measurements of model performance. The first, which we call accuracy, is simply the proportion of individuals in a cluster for which the infector with the highest posterior support was in fact the true infector. For some tests, we also used the mean of the posterior support values for the true infector, which we call the true posterior probability.

### Effect of Biomarker Information

To test the potential of improved transmission history inference using real biomarkers, we used a transmission tree with fixed infection times and 15 individuals, mean temporal spacing of 0.5 years, and transmission heterogeneity of about 1.24, generating 100 different transmission histories per tree with different sets of sampling times for each history. For each individual in each history, the time between infection and sampling is determined by independent Gamma distributed random variables with mean 2 years and standard deviation 1.5 years. Sequences were generated for each phylogeny using both the *env* and *pol* mutation parameters. We simulated biomarker values for all of the infection ages at the time of sampling, then ran the multiple biomarker model to infer the infection time distributions. In order to assess the effect of the amount of biomarker information, we ran transmission history inference with the infection age distributions using 2, 3, or 5 biomarkers as well as infection age distributions representing no or uninformative biomarkers and near perfect infection age information that effectively provided fixed infection times. In the no biomarker case, the infection age distribution was a continuous uniform distribution with minimum 0 and maximum 11 years. In the fixed infection time case we used a Gamma distribution with mean equal to the true infection age and standard deviation equal to 0.005 years.

### Effects of Transmission Cluster Attributes

To test the effect of various attributes of the transmission cluster itself, one variable at a time, we investigated 1) the number of individuals, ranging from 5 to 50, 2) the temporal spacing, ranging from 0.01 years to 2.5 years, and 3) the transmission heterogeneity, ranging from 0 to 3.74 (the maximum possible for 15 individual clusters). While each attribute was varied, the other attributes were held constant, with the non-variable values at 15 individuals, temporal spacing of 0.5 years, and transmission heterogeneity around 1 (between 0.8 and 1.2). Both the *env* and *pol* mutation models were used for each history. In all trials, we used simulated infection age distributions using all five biomarkers. We used two subsets of trials, one with more realistic HIV values and one corresponding to an idealized situation. For the realistic values, we again used *α* = 5 and *β* = 5 for the within-host model, two years between infection and sampling, and independent biomarkers for each individual in a given transmission cluster. For the idealized situation, we used *α* = 0 and *β* = 0.1 (resulting in a short pre-transmission interval [12, 18] where the phylogeny closely resembles the transmission tree), one year between infection and sampling, and fixed biomarkers for all individuals.

We also tested how different attributes of the transmission clusters may interact with each other, possibly affecting the difficulty of inference. To do this, we simulated histories with all combinations of different values for each attribute. These simulated clusters had 10, 15, 20, or 40 individuals, temporal spacing of 0.1, 0.5, 1, or 2 years, transmission heterogeneity near 0, 0.5, 1, 1.7, 2.3, or 3, with both substitution models (*env* and *pol*) and all five levels of biomarker information used with each cluster.

### Effect of Multiple Sequences per Individual

To test whether additional sequence data per patient can help counteract the inference problems inherent with wide transmission bottlenecks, we simulated 200 transmission histories with 3 individuals with temporal spacing of 0.5 years, including both the serial infection case and the case where the first individual infects the other two. We simulated phylogenies on each transmission history with 4 sampled sequences per individual taken two years after the time of infection, and *α* = 5, *β* = 5. Next, we generated subsampled phylogenies, keeping only 1 sequence per individual for each tree. Finally, inference was performed on both the full and subsampled datasets using 1×10^6^ iterations of MCMC after 4×10^4^ iterations of burn-in, with some runs concluding sooner if the target ESS is reached early.

### Effect of Incomplete Sampling of Transmission Clusters

To assess the impact of violating the assumption of complete sampling, we ran the inference method on both complete and incomplete simulated transmission histories. We used four different basic transmission histories with the amount of transmission heterogeneity varying from none to moderately high, with the infectors, infection times, and sampling times fixed within each of the four trees, while the phylogenies and sequences for each replicate were generated independently. The phylogenies were generated using the coalescent simulation with two different values of *α*, corresponding to wide (*α* = 5) and complete (*α* = 0) transmission bottlenecks, while *β* remained at 5. In each case, transmission history inference was performed on both the complete transmission history as well as that same transmission history with the data from the individual that infected the most other individuals removed (in the no transmission heterogeneity case, the sixth individual is removed). In both cases, there was no penalty for transmission after diagnosis. Accuracy is defined as before except that in the incomplete sampling case, we considered the inference to be “accurate” when the true infector’s infector is chosen when the true infector is not sampled. In addition to the overall accuracy, we also looked at the accuracy of the individuals whose true infector is not sampled on their own.

### Real HIV Transmission Cluster Data

We demonstrate the application of the inference method on data from 4 real transmission clusters involving 4-14 patients in the larger Swedish HIV epidemic that are believed to be at least close to fully sampled [17, 38, 39]. These data included sequence data from *pol* drug resistance testing, biomarkers BED, CD4, and *pol* polymorphisms, as well as first positive test dates for all patients, and some patients had a previous negative test date.

We first used the MBM to infer the distributions of the infection times for all individuals. The previous negative tests were accounted for by assuming that individuals could not have been infected more than two months before the most recent negative test, then scaled the prior distribution between earliest possible infection time and the diagnosis date, taking into account that patients without regularly scheduled tests are typically infected closer to the first positive test than the last negative test [28]. With the numeric distributions for the infection times, we used *biophybreak* with 2×10^6^ MCMC iterations (5×10^4^ iterations of burn-in) on each transmission cluster.

## RESULTS

### An improved multi-biomarker model for estimation of HIV-1 time of transmission

Using the 30-patient training data, we modeled five biomarkers as linear-asymptotic trends for BED and LAg, and linear for *pol* polymorphism count, *pol* NGS diversity, and CD4 cell count (Fig 2). The biomarkers were combined into a mixed effects modeling framework to allow for patient specific variation and general trends. As expected, a shorter time between infection and sampling typically resulted in a posterior distribution with lower standard deviations, and longer time between infection and sampling resulted in more uncertainty about the time of infection.

Including all five biomarkers, we were able to substantially improve the performance over the previous 3-biomarker model (*pol* polymorphisms + CD4 + BED) used in Giardina et al [21], with more than 2-fold reductions in mean bias, mean absolute error (MAE), and root mean square error (RMSE) when comparing the medians of the inferred distributions in a cross-validation to the true values (Table 1). We also evaluated the performance of our modified 3-biomarker model (*pol* polymorphisms + CD4 + BED) as well as a 2-biomarker model (*pol* polymorphisms + CD4), which is of practical interest because *pol* polymorphism count and CD4 cell counts almost always exist in HIV-1 clinical databases. Our new 2- and 3-biomarker models also improved over the previous 3-biomarker model.

**Table 1.** Biomarker model performance.

| Model | Mean Bias | MAE | RMSE |
| --- | --- | --- | --- |
| 3 Biomarkers, 2019 | -0.68 | 1.01 | 1.38 |
| 5 Biomarkers, 2021 | -0.19 | 0.33 | 0.47 |
| 3 Biomarkers, 2021 | -0.23 | 0.43 | 0.58 |
| 2 Biomarkers, 2021 | -0.36 | 0.49 | 0.70 |

Table Footnote: All values are in years relative to actual time of infection. The 2019 model is from Giardina et al [21], shown for comparison; the 2021 models are those developed in this study. 5 biomarkers = BED, LAg, *pol* polymorphisms, *pol* NGS diversity, and CD4 cell count; 3 biomarkers = BED, *pol* polymorphisms, and CD4 cell count; 2 biomarkers = *pol* polymorphisms and CD4 cell count.

### Biomarker information significantly improves transmission reconstruction

To investigate the accuracy of adding different numbers of biomarkers to a coalescent-based transmission model, we investigated 1,000 simulations with varying times between infection and sampling, and sampling different, possible phylogenies on a fixed transmission tree with moderate values for transmission heterogeneity and temporal spacing (Fig 3). In these simulations we used only one sequence per host as that is the standard in clinical and public health databases. Note, however, that our phylogenetic framework models within-host diversity, which can be seen in the reconstructions involving multiple transmitted lineages and super-spreader activity.

**Fig 3:**
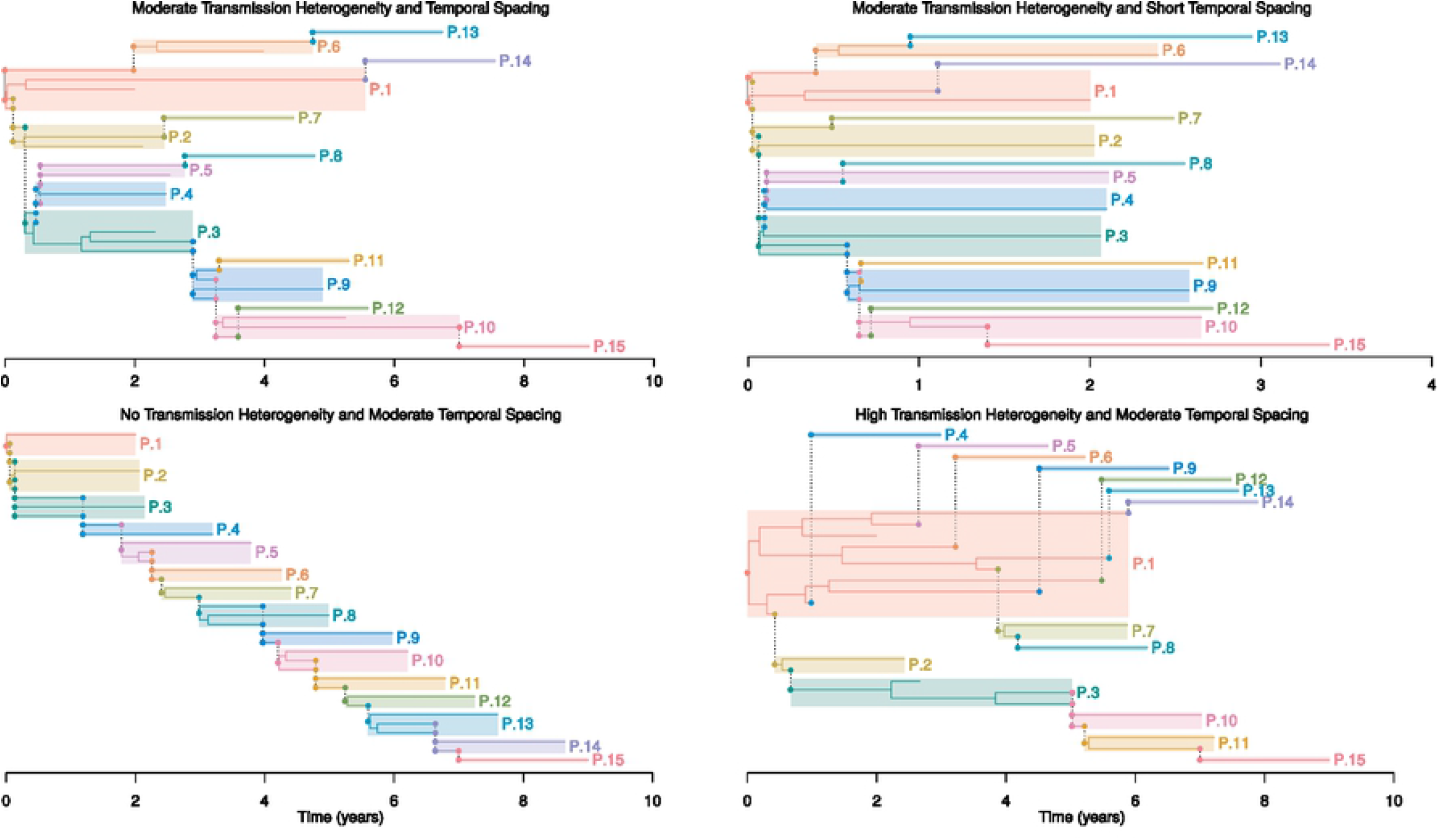
Example Transmission Histories. Four different possible transmission histories and phylogenies. As in Fig 1, each host is indicated by a colored box and transmissions are indicated by dashed vertical lines. Because transmission after diagnosis is not prohibited in these examples, it is possible that the right side of the box for some individuals to extend beyond the sampling time, in which case the right side of the box is the time of the last transmission and the sampling time is the point of termination of a lineage in the interior of the box. Note also that although we only sample one lineage here, the model takes within-host diversity into account, and thus can infer which lineage(s) within the host’s diversity that was transmitted. The Top Left Panel shows a transmission history with moderate temporal spacing (0.5 years) and transmission heterogeneity (near 1). The Top Right Panel shows a transmission history with moderate transmission heterogeneity (near 1) and short temporal spacing (0.1 years). The Bottom Left Panel shows a transmission history with no transmission heterogeneity, leading to a straight transmission chain. The Bottom Right Panel shows a transmission history with high transmission heterogeneity involving a super-spreader (P.1).

Adding real biomarker information about time of infection significantly improves the accuracy in reconstructing transmission histories (Fig 4). We compared adding 2, 3, or 5 biomarkers, as well as fixed infection times, to phylogenetic information only from HIV-1 transmissions in a coalescent-based modeling system. This is a non-trivial problem because virus phylogenies from epidemiologically linked patients cannot be assumed to be identical to the transmission history among the patients [12, 17]. Here, we investigated the overall expected probability to infer the correct donor in each transmission among 15 patients. While the use of sequence data and phylogenetic reconstruction is much better than a random guess at 1/N, increasing from 6.7% expected accuracy to 30% with no biomarkers, we improved the transmission history inference over the phylogeny alone by on average 12 percentage points using the broadly available 2 biomarkers *pol* polymorphisms and CD4 cell counts (p < 1×10^-9^, Wilcoxon signed rank test with Bonferroni multiple testing correction). Adding the 3-biomarker model improved the accuracy by >13 percentage points (p < 4×10^-10^) and adding all 5 biomarkers by 16 percentage points (p < 4×10^-12^). We investigated the theoretical limit of using biomarker data to our transmission history inference by adding effectively fixed infection times, which reached on average a 29 percentage point accuracy improvement over the phylogeny alone. All improvements in accuracy were achieved by an increase in the model posterior prediction score (Fig S1).

**Fig 4:**
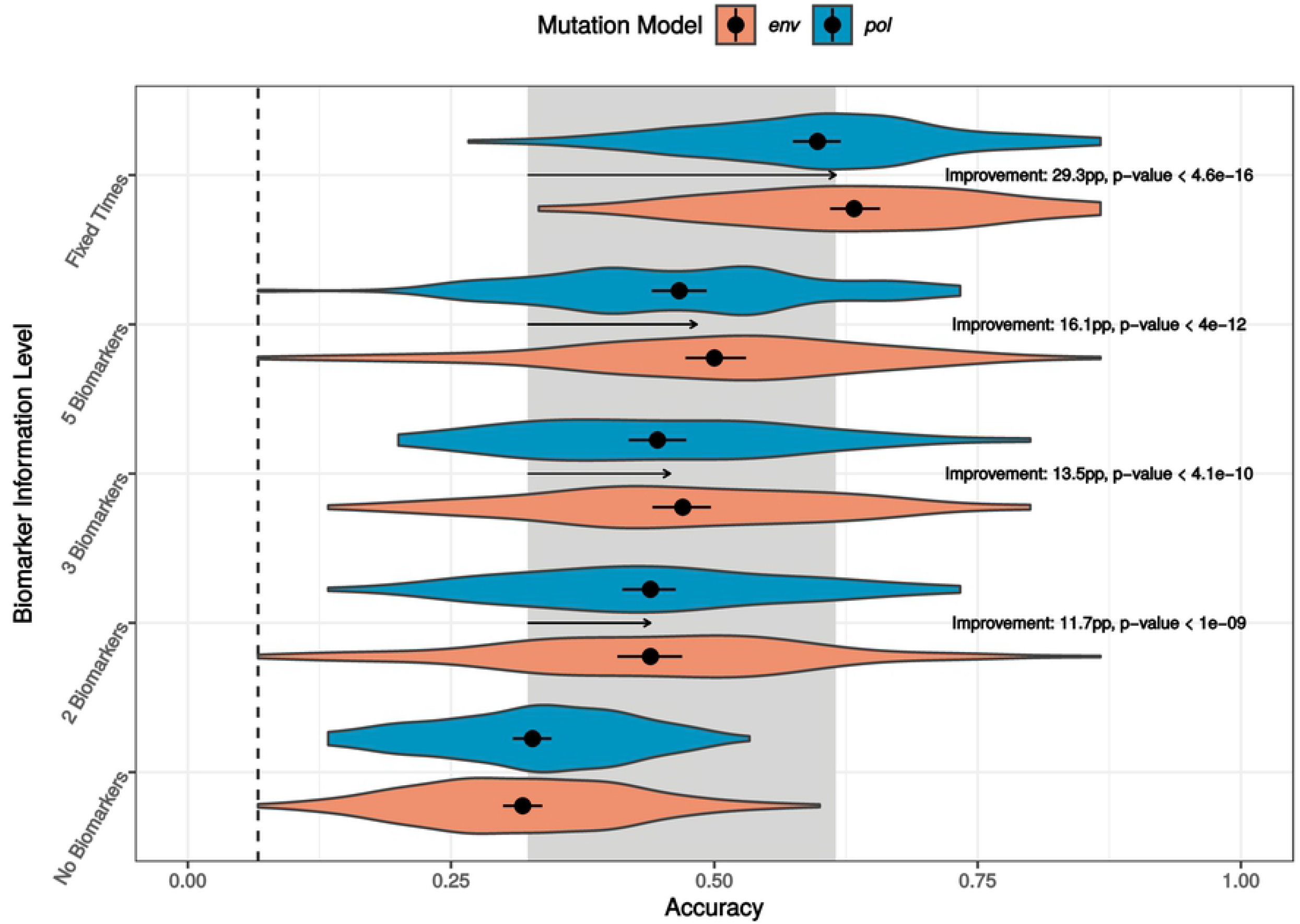
Transmission Inference Improvement with Biomarker Information. Violins show the full distribution of accuracy on simulated data for each level of biomarker information and mutation model (genomic region), with the point and line segment in each violin representing the mean and 95% bootstrapped interval for the estimate of the mean. The gray shaded area in the background represents the region between effectively no information about infection times and effectively fixed infection times, with the information level attainable with biomarkers falling between these two extremes. The improvement is shown with an arrow in percentage points (pp) and p value estimated by a Wilcoxon signed rank test with Bonferroni multiple testing correction.

Overall, the *env* gene performed somewhat better than *pol* in the combined biomarker-phylogenetic inference of the transmission history. Part of the explanation likely lies in the fact that *env* evolves faster, thus accumulating more information about genealogical relationships, making the phylogenetic component more robust as previously shown [40, 41].

### Shorter temporal spacing and increased transmission heterogeneity reduce reconstruction accuracy, but larger clusters are not harder to get right

Heterogeneity in the number of transmitted lineages (phylogenetically separate virus variants), time between infection and onward transmission, time between infection and sampling, and in the number of onward infections a host causes (transmission degree) are all known to occur in real transmission histories. Therefore, using 5 biomarkers, we modeled all of these factors, as well as different sizes of transmission clusters, and assess their effects on the accuracy of transmission history inference. These scenarios cover a wide range of fully sampled, possible, realistic HIV-1 transmission histories (Fig 3).

Under realistic HIV-1 evolutionary within-host parameters (*α* = 5, *β* = 5), where about half of transmissions result in >1 transmitted lineage and within-host diversification is substantial [30], transmission history inference is expected to be quite challenging. For comparison, if only single lineages were transmitted and within-host diversification was very limited (*α* = 0, *β* = 0.1), the overall accuracy reaches about 90% when infections were not close in time and transmission degree was 1 or less (Fig 5A). With realistic HIV-1 parameters, the overall accuracy was about 50% (Fig 5B).

**Fig 5:**
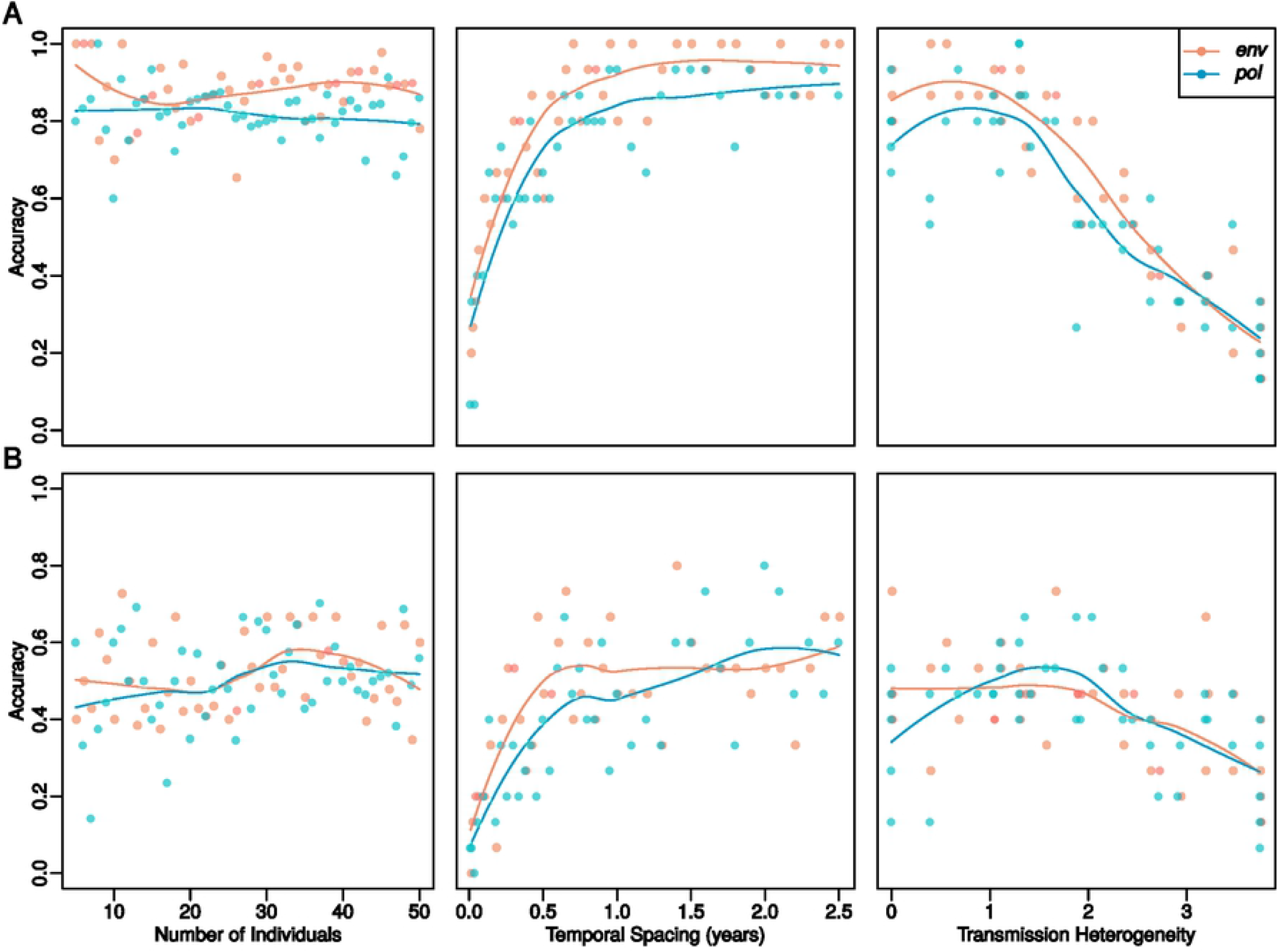
Individual Transmission Cluster Attribute Effects. Each point represents the accuracy of inference on a single transmission history, with the solid lines being the loess lines for each mutation model (*env* or *pol* genomic regions). (A) The top row shows simulation results from idealized situations with complete transmission bottlenecks, low within host diversity, one year between infection and sampling, and fixed biomarkers. (B) The bottom row shows results from simulations with realistic HIV parameters in terms of transmission bottleneck size variation and within host diversity, two years between infection and sampling, and independent biomarkers.

The shorter the temporal spacing of transmission events, the harder it was to infer the correct transmission history. Although biomarkers improved the reconstruction accuracy even at very short temporal spacing of only a few days, raising the accuracy over a random guess at 1/N, meaningful accuracy started when the temporal spacing was above a few months. This is because biomarker posterior distributions will greatly overlap when times between transmission are short, making it difficult to order the events in time. With short times between many infections, there was also little time to accumulate mutations that would inform the phylogenetic reconstruction.

Higher degrees of transmission heterogeneity, like in panel 4 in Fig 3, on average also lead to more difficult transmission history inference. At degree levels above 2.5, the average accuracy decreased from about 50% to 25%. Compared to the overall performances and limits in Fig 4, this reduction was as severe as not having any biomarkers, and clearly constitutes a very difficult to resolve epidemiological situation. The most difficult cases were those where there were short temporal spacing and high transmission heterogeneity. Thus, super-spreader activity can cause particularly difficult to reconstruct epidemiological scenarios where biomarkers may not always help to resolve the transmission history.

Most combinations of attributes showed only small amounts of interaction effects. The most notable exception was the temporal spacing and biomarker information level, which had a combined effect on the quality of inference when marginalizing over the other three variables (Fig S2). With short temporal spacing, the biomarker information offered only small improvements. Having near perfect information about the infection times (the fixed infection time case), however, would allow a very large improvement. As the temporal spacing increases, the improvement with better biomarker information increased as well, while also approaching the fixed infection time case. This is because as the temporal spacing increases, the infection time distributions become more separated, allowing greater certainty about the infection order. In this way, longer temporal spacing helps both in terms of making the phylogeny more informative and helping the biomarkers to allow more separation.

Promisingly, transmission histories involving more hosts were on average not harder to reconstruct at fixed levels of transmission heterogeneity and temporal spacing (Fig 5). This is encouraging for real-time applications that follow the growth of a public health database because it cannot be known beforehand how many persons that eventually will be part of a transmission cluster. Also encouraging was that *env* and *pol* performed similar in these simulations, as public health databases typically store *pol*, but not *env*, sequences from drug resistance testing.

### Additional sequences from hosts improve overall transmission reconstruction

Beyond simply having more data, the conceptual motivation for using >1 sequence/host is that it should increase the chance that the sampled lineages from a recipient will coalesce with at least one of the sampled lineages from the donor rather than earlier in the transmission history (Fig S3). Although the computational burden increased when adding more sequence data per infected host, the accuracy in the transmission history reconstruction did indeed improve. Using 4 sequences instead of 1 sequence per host in 3-person transmission histories showed a 7.4 percentage point (12.2%) improvement in accuracy (on average 0.073 posterior probability (14.9%) improvement; p < 2.5×10^-7^, Wilcoxon signed rank test) (Fig S4). While encouraging for future analyses with richer sequence data, further development of computational efficiency will be needed to exploit this enhancement.

### The overall accuracy is not significantly affected by incomplete sampling

While the “first” person’s donor will always be missing, and recipients that have not infected anybody may also be irrelevant to the transmission-history-reconstruction-problem, the problem of missing intermediary links is always a possibility. Thus, in real-life situations it is never known if the sample is complete or not, i.e., one cannot be sure that all *relevant* donors have been sampled. Therefore, we investigated the situation when an intermediary donor was missing in transmission histories with 10 hosts, using trees with four levels of transmission heterogeneity (Fig S5). When missing, we defined accurate donor identification as the missing donor’s donor.

The differences in accuracy between the completely and incompletely sampled transmission histories were in general relatively small (Fig 6). In terms of the overall accuracy of the inference, the absolute difference in mean accuracy between the completely and incompletely sampled transmission clusters was less than 2.6 percentage points for any combination of bottleneck size and transmission heterogeneity (all raw p-values > 0.05, Wilcoxon signed-rank tests).

**Fig 6:**
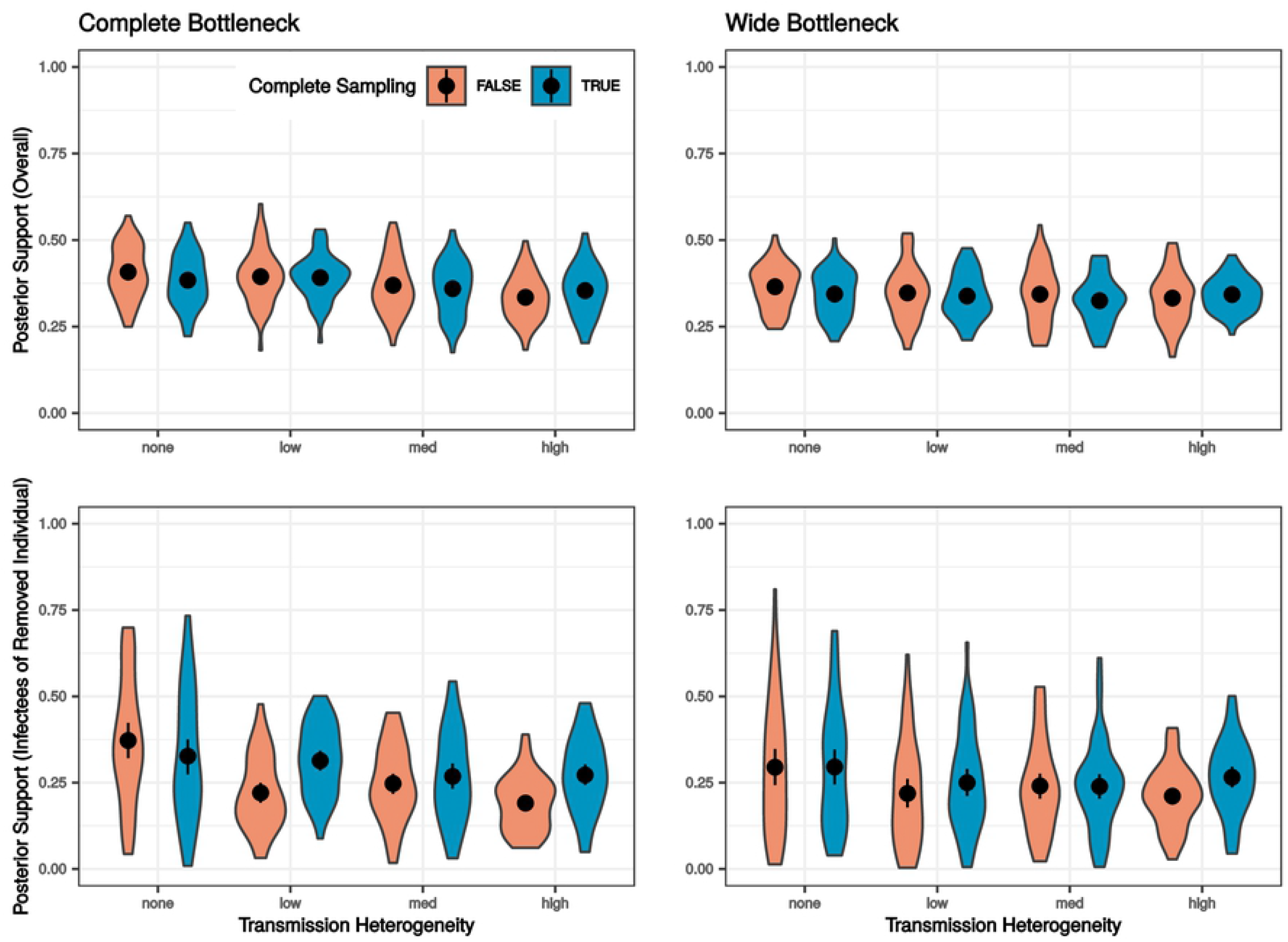
The Effect of Incomplete Sampling. Violins show the distribution of means of the posterior probability for true infectors (donors), or the true infectors’ infectors if the true infector is unsampled, with the point and line segment in each violin representing the mean and 95% bootstrapped interval for the estimate of the mean. (Top Row) Overall model performance. (Bottom Row) Model performance for the individuals infected by the unsampled individual.

Focusing on the accuracy of assigning the donor to the recipients infected by the unsampled donor, naturally, was the most challenging. Three out of the eight combinations of bottleneck size and transmission heterogeneity showed moderate drops in accuracy, as well as significant p-values while the other five had only small and insignificant differences. The three situations with moderate differences were the low and high transmission heterogeneity cases with complete bottlenecks and the high transmission heterogeneity case with the wide bottleneck, with the differences in accuracy at 20.0, 11.2, and 8.4 percentage points, respectively (raw p-values at 0.002, 0.015, and 0.046, Wilcoxon signed-rank test).

Although there were some cases where the accuracy of inference for individuals infected by an unsampled individual was lower, since most differences are small and the overall accuracy remained at the same level, these results demonstrate that it is appropriate to apply this method even when the assumption of fully sampled transmission clusters is violated.

### Application to real transmission clusters

To assess transmission history reconstruction using real data, we applied our method to four transmission clusters from the general Swedish HIV-1 epidemic [17, 38, 39]. The data included direct population *pol* gene sequences [25], determined as part of clinical drug resistance testing, BED and CD4 T cell counts, occasionally previous negative tests, and date of sampling. The sequence data was used for phylogenetic inference as well as a biomarker of within-host divergence (*pol* polymorphisms). Fig 7 shows an inferred chain of four sampled hosts infecting each other serially (Fig 7A) and a more complex transmission history involving 13 sampled hosts that included super-spreading (Fig 7B). When longer time from infection to transmission occured and the biomarker density was narrow, the posterior support for donor assignment was high, e.g., the donor of P.85.1480 is assigned to P.85.1368 at 0.99 posterior support, who transmitted (at least) two HIV-1 lineages (Fig 7A). Conversely, when there were short time intervals between transmissions and biomarker densities overlap, transmission reconstruction became more difficult, e.g., the donor of P.85.1173 was more evenly attributed to 3 out of 4 sampled hosts in the corresponding transmission cluster (Fig 7A), and, similarly, assigning a donor to P.24.323 was less certain in the larger cluster (Fig 7B). Because phylogeny, biomarkers, and sampling times interact in non-trivial ways, however, relatively large posterior probabilities may be assigned to one donor over many others, e.g., in the transmission to P.24.859, P.24.909 was significantly more likely to be the donor than any other sampled donor in that cluster (Fig 7B). Fig S6 shows two additional transmission clusters with transmission heterogeneity (degrees 1 and 1.5), overlapping biomarkers, and both short overall time (6 transmissions in <1 year) and longer time (8 transmissions over 8 years). These clusters provide further examples of real situations where some donor assignments were easier and others harder.

**Fig 7:**
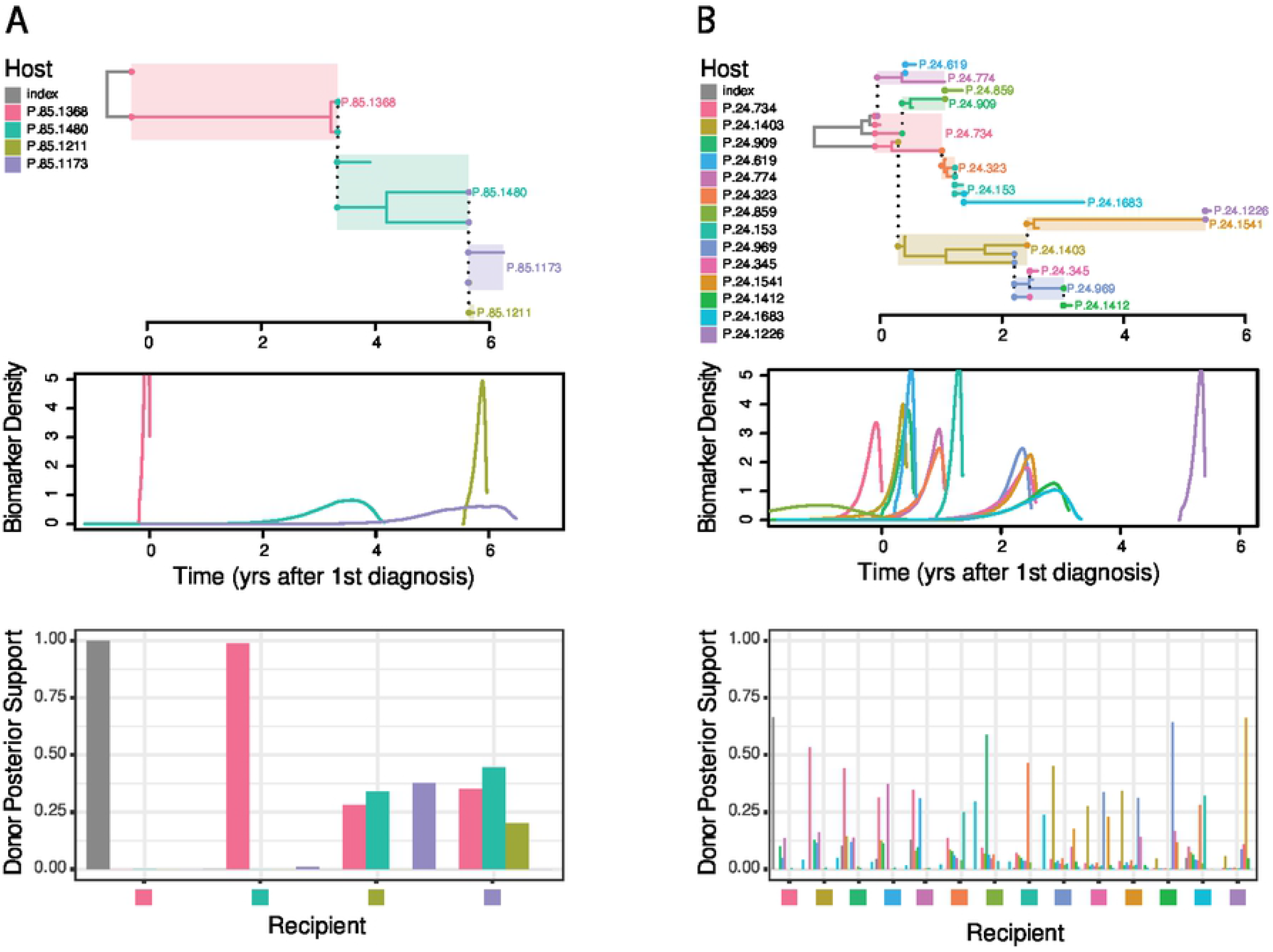
Transmission History Inference on Real HIV Transmission Clusters. Examples of a smaller, simple transmission history (A) and a larger, complex transmission history (B) from the Swedish HIV epidemic. The Top Panel in each transmission cluster shows the inferred maximum parent credibility tree. The Middle Panel shows the distributions of infection times inferred from biomarker values for each individual, using our 3-biomarker model applied to data that existed in a public health database. The Bottom Panel shows posterior support for each individual to be the donor for each individual, with each colored square on the x-axis representing one individual and the height of the colored bars represent the posterior support for the corresponding individual to be their infector.

## DISCUSSION

In this study we have developed a biomarker-enhanced phylogenetic framework to allow for more accurate inference of pathogen transmission histories. The biomarker component used five real HIV-1 biomarkers from a set of untreated, longitudinally followed HIV-1 infected patients. Thus, our results display practical and realistic improvement of the expected accuracy in the inference of HIV-1 transmission histories using data that is typically available in a HIV surveillance system. We investigated a wide variety of transmission scenarios, including heterogeneity in the number of transmitted lineages, in the time between infection and onward transmission, in the time between infection and sampling, in the number of onward infections a host causes, and among different sizes of transmission clusters. Overall, we show that adding biomarkers to the transmission history inference substantially improved accuracy in all the considered scenarios.

Compared to previous phylodynamic methods that infer transmission history and direction [4–7, 10], our method includes real biomarker-informed time of transmission and allows for wide transmission bottlenecks. Kenah et al previously showed that if the relative order of transmissions is known, transmission history inference should improve [20]. Here, we show that real HIV biomarkers approach this ideal situation, where using more biomarkers is better than fewer, but it is unlikely that we will ever find biomarkers that can resolve all situations. Likewise, factors such as within-host diversity of the virus, the fact that HIV transmission largely is a random draw of a few variants [17], transmission often involves >1 phylogenetic lineage [30], and that within-host evolution involves lineage death and birth [12, 42, 43], together put theoretical limits on how accurately we can infer the underlying transmission history from a phylogeny.

While our biomarker-enhanced phylogenetic method generally improved transmission history inference, one should not expect such a method to reach 100% accuracy. Here, we show that even if the biomarkers could provide perfect transmission times, having realistic levels of within-host virus diversity induces substantial uncertainty that limits the average possible performance to 40-87% accuracy (95% posterior probability interval) assuming moderate temporal spacing and transmission heterogeneity. There are also certain situations that are particularly difficult to accurately reconstruct. When transmission histories involve large transmission heterogeneity, typically when super-spreader activity has occurred, it becomes difficult to reconstruct all transmission events accurately. This is in part because each time the super-spreader transmits, a random draw of variants is transmitted and thus the phylogenetic ordering of coalescences typically does not follow the transmission order [12, 17]. Furthermore, two hosts infected close in time to each other may receive more similar virus than is later sampled in the donor, and thus may appear to be linked to each other rather than to the donor. This complication is compounded when many transmissions happened over a short time. It is possible that prior identification of super-spreader activity can identify when and where in a phylogeny these problems exist [44].

Another uncertainty is related to the fact that we never have a perfect sample from an ongoing epidemic, meaning that, at any time, we have not yet sampled every actual or soon to be donor. This is highlighted by the fact that many populations still are far from the WHO/UNAIDS 90-90-90 goal [45, 46], and even in nations where that goal has been reached it takes on average 2 years to detect most infections [47]. However, we show that our biomarker enhanced phylogenetic framework can handle missing links quite well, typically identifying a missing donor’s donor as the source of the transmission. Also, note that missing individuals that have not infected anyone else leave no trace in a phylogeny. Therefore, a missing link refers to any unsampled person ancestral to the set of sampled persons. While the existence of missing individuals who have not infected any sampled individuals is certainly a concern from a public health perspective, this is not something that can be determined from the available sequence or biomarker data without making substantial assumptions about the expected number of new individuals infected per host.

Phylogenetic reconstruction of transmission histories is a powerful and scientifically sound method because it is objective, can evaluate alternative hypotheses, and, as we show here, can be augmented with additional data. Because HIV infection still causes stigma and legal risks in some jurisdictions, however, both research and public health projects that use such methodology must be ethically justified on the basis of providing public health benefits [48, 49]. Here, we show that phylogenetic methods can be made more accurate by adding biomarker data on time of infection. Accuracy is important because it allows public health resources to be directed to where they are needed most, and thus will have the largest reduction in disease spread [50]. Again, we emphasize that it can never reach 100% certainty, typically much less, yet the levels we can reach with the proposed methodology should make public health efforts more efficient.

Improved HIV surveillance, source attribution, and outbreak response depends on advances in HIV prevention, diagnosis, and continuous treatment. Application and further development of the technology presented here could allow for better prevention programs focusing on locally informed and tailored strategies.

## ACKNOWLEDGEMENTS

We are grateful for helpful discussions with Dr. Don Klinkenberg, and for his unpublished modifications to the R package *phybreak* that allow for wide transmission bottlenecks. We are also grateful to the HIV infected study participants and to Johanna Brodin, Fabio Zanini, Lina Thebo, Eva Eriksson, Christa Lanz, and Göran Bratt for important contributions in the generation of the published data that was analyzed in this study. We thank Helena Skar, Federica Giardina, Tom Britton, Vadim Puller, and Richard Neher for previous developments of the theoretical models of biomarker-based estimation of HIV infection times. This study was supported by NIH/NIAID grants R01AI087520 and R01AI152897.

## FIGURE LEGENDS

**Fig S1 Relationship Between Accuracy and True Posterior Probability**

Accuracy and mean true posterior probability values for each of the 1000 trials used while testing the effect of biomarker information shown with the *y* = *x* line for comparison. Note that since all trials are on clusters with 15 individuals, the accuracy for each trial can only be one of sixteen possible values, while the means of the true posterior probability are effectively continuous.

**Fig S2: Combined Transmission Cluster Attribute Effects**

Violins show the distribution of accuracy for each level of biomarker information for each amount of temporal spacing, with the point and line segment in each violin representing the mean and 95% bootstrapped interval for the estimate of the mean.

**Fig S3: Conceptual Motivation for Additional Sequences per Individual**

(Top) A transmission history with 4 sampled sequences per individual. (Bottom) The same history subsampled to 1 sequence per individual.

**Fig S4: Inference with Multiple Sequences**

Violins show the distribution of mean true posterior support for 4 and 1 sequence(s) per individual for each mutation model, with the point and line segment in each violin representing the mean and 95% bootstrapped interval for the estimate of the mean.

**Fig S5: Transmission Histories used in Incomplete Sampling Tests**

We test the effect of an unsampled individual with four different levels of transmission heterogeneity. The 6^th^ individual is removed in the cases of no and low transmission heterogeneity, the 5^th^ individual is removed in the case of moderate transmission heterogeneity, and the 3^rd^ individual is removed in the case of high transmission heterogeneity.

**Fig S6: Additional Real Transmission Clusters**

(Top) Inferred maximum parent credibility trees for each transmission cluster. (Middle) Distributions of infection times inferred from biomarker values for each individual. (Bottom) Posterior support for each individual to be the donor for each individual, with the height of the colored bars represent the posterior support for the corresponding individual to be their infector.

